# *Candidatus* Nitrosopolaris, a genus of putative ammonia-oxidizing archaea with a polar/alpine distribution

**DOI:** 10.1101/2021.12.10.472193

**Authors:** Igor S. Pessi, Aino Rutanen, Jenni Hultman

## Abstract

Ammonia-oxidizing archaea (AOA) are key players in the nitrogen cycle of polar soils. Here, we analysed metagenomic data from tundra soils in Rásttigáisá, Norway, and recovered four metagenome-assembled genomes (MAGs) assigned to the genus “UBA10452”, an uncultured lineage of putative AOA in the order Nitrososphaerales (“terrestrial group I.1b”), phylum Thaumarchaeota. Analysis of other eight previously reported MAGs and publicly available amplicon sequencing data revealed that the UBA10452 lineage is predominantly found in acidic polar and alpine soils. In particular, UBA10452 MAGs were more abundant in highly oligotrophic environments such as mineral permafrost than in more nutrient-rich, vegetated tundra soils. UBA10452 MAGs harbour multiple copies of genes related to cold tolerance, particularly genes involved in DNA replication and repair. Based on the phylogenetic, biogeographical, and ecological characteristics of 12 UBA10452 MAGs, which include a high-quality MAG (90.8% complete, 3.9% redundant) with a nearly complete 16S rRNA gene, we propose a novel *Candidatus* genus, *Ca*. Nitrosopolaris, with four species representing clear biogeographical/habitat clusters.

## Introduction

Nitrification – the oxidation of ammonia to nitrite and further oxidation to nitrate – is a crucial part of the nitrogen (N) cycle providing a link between reduced and oxidized forms of N. The first step of nitrification, ammonia oxidation, is carried out mainly by aerobic chemolithoautotrophic microorganisms that grow by coupling the energy obtained from the oxidation of ammonia with carbon dioxide (CO_2_) fixation (Lehtovirta-Morley, 2018). Ammonia-oxidizing archaea (AOA) outnumber ammonia-oxidizing bacteria (AOB) by orders of magnitude in many terrestrial and aquatic environments, particularly in oligotrophic environments with low N input (Leininger *et al*., 2006; Schleper and Nicol, 2010; Lehtovirta-Morley, 2018). Among the reasons for their ecological success is an enzymatic machinery with higher affinity for ammonia and a more efficient CO_2_ fixation pathway than their bacterial counterparts (Martens-Habbena *et al*., 2009; Könneke *et al*., 2014; Kerou *et al*., 2016). However, high ammonia affinity is not a common trait to all AOA, with some strains displaying a low substrate affinity that is comparable to that of non-oligotrophic AOB (Kits *et al*., 2017; Jung *et al*., 2022).

Ammonia oxidation is an important process in polar soils despite commonly N limited and cold conditions (Alves *et al*., 2013; Siljanen *et al*., 2019; Hayashi *et al*., 2020). AOA generally outnumber AOB in oligotrophic polar soils and are often represented by few species (Alves *et al*., 2013; Magalhães *et al*., 2014; Richter *et al*., 2014; Pessi *et al*., 2015, 2022, pre-print; Siljanen *et al*., 2019; Ortiz *et al*., 2020). Due to their predominance, AOA are important contributors to the N cycle in polar soils and are thus key players in the cycling of the potent greenhouse gas nitrous oxide (N_2_O). Contrary to earlier assumptions, polar soils are increasingly recognized as important sources of N_2_O (Voigt *et al*., 2020). Both the nitrite originated from the oxidation of ammonia as well as the nitrate produced in the second step of nitrification are the substrates of denitrification, an anaerobic process that has N_2_O as a gaseous intermediate (Butterbach-Bahl *et al*., 2013). Moreover, AOA have been directly implicated in the production of N_2_O under oxic conditions via several mechanisms such as hydroxylalamine oxidation and nitrifier denitrification (Wu *et al*., 2020). However, both the direct and indirect role of AOA in the cycling of N_2_O is much less understood compared to their bacterial counterparts.

AOA are notoriously difficult to cultivate and so far only three genera have been formally described based on axenic cultures: *Nitrosopumilus* (Qin *et al*., 2017) and *Nitrosarchaeum* (Jung *et al*., 2018) in the order Nitrosopumilales (“marine group I.1a”) and *Nitrososphaera* (Stieglmeier *et al*., 2014) in the order Nitrososphaerales (“terrestrial group I.1b”). Several provisional *Candidatus* genera have also been proposed based on non-axenic enrichments, e.g. *Ca*. Nitrosocaldus (“termophilic group”) (de la Torre *et al*., 2008) and *Ca*. Nitrosotalea (“group I.1a-associated”) (Lehtovirta-Morley *et al*., 2011). Moreover, the growing use of genome-resolved metagenomics has resulted in the identification of tens of novel, currently uncultured lineages in the phylum Thaumarchaeota (Rinke *et al*., 2021). These lineages are phylogenetically distinct from both formally described and *Candidatus* taxa and are identified with placeholder alphanumeric identifiers (e.g., the Nitrososphaerales genus “UBA10452”). The identification of these novel lineages by metagenomics greatly expands our knowledge of the diversity of AOA but detailed descriptions of their metabolic and ecological features are generally lacking.

Recently, we have applied a genome-resolved metagenomics approach to gain insights into the microorganisms involved with the cycling of greenhouse gases in tundra soils from Kilpisjärvi, Finland (Pessi *et al*., 2022, pre-print). Analysis of *amoA* genes encoding the alpha subunit of the enzyme ammonia monooxygenase (Amo) revealed a very low diversity of ammonia oxidizers, with only four genes annotated as *amoA* out of 23.5 million assembled genes. Three of these were most closely related to the *amoA* gene of the comammox bacterium *Ca*. Nitrospira inopinata (Daims *et al*., 2015). The remaining *amoA* gene was binned into a metagenome-assembled genome (MAG) assigned to the genus “UBA10452”, an uncharacterized archaeal lineage in the order Nitrososphaerales, phylum Thaumarchaeota (Rinke *et al*., 2021). Here, we i) report four novel UBA10452 MAGs obtained from tundra soils in Rásttigáisá, Norway; ii) characterize the genomic properties, metabolic potential, phylogeny, and biogeography of the UBA10452 lineage; and iii) propose the creation of a new *Candidatus* genus, *Ca*. Nitrosopolaris.

## Methods

### Sampling and metagenome sequencing

Ten soil samples were obtained in July 2017 across an area of alpine tundra in Rásttigáisá, Norway (69°59’N, 26°15’E, 700 m.a.s.l.). DNA was extracted from the mineral layer (10–15 cm depth) with the PowerSoil DNA Isolation kit (QIAGEN, Venlo, Netherlands) according to the manufacturer’s instructions. Paired-end metagenomic sequencing was done using the Illumina NextSeq500 platform (Illumina, San Diego, CA, USA) at the DNA Sequencing and Genomics Laboratory (Institute of Biotechnology, University of Helsinki).

### Metagenome assembling and binning

Removal of adapter sequences and low-quality base calls (Phred score < 28) was done with Cutadapt v1.10 (Martin, 2011) and sequences were assembled with MEGAHIT v1.1.1 setting a minimum contig length of 1,000 bp (Li *et al*., 2015). Samples were assembled individually and as one co-assembly of all samples pooled together. Manual MAG binning was done with anvi’o v6.0 (Eren *et al*., 2015) according to Pessi *et al*. (2022, pre-print). In brief, Prodigal v2.6.3 (Hyatt *et al*., 2010) was used to predict gene calls and single-copy genes were identified with HMMER v.3.2.1 (Eddy, 2011). Bowtie v2.3.5 (Langmead and Salzberg, 2012) and SAMtools v1.9 (Li *et al*., 2009) were used to map the quality-filtered Illumina reads to the contigs. Contigs were then manually binned into MAGs based on differential coverage and tetranucleotide frequency using the *anvi-interactive* interface of anvi’o v6.0. MAGs were manually inspected and refined using the *anvi-refine* interface of anvi’o v6.0.

### Metagenome-assembled genomes assigned to the UBA14052 lineage

MAGs were classified based on 122 archaeal and 120 bacterial single-copy genes with GTDB-Tk v1.3.0 (Chaumeil *et al*., 2019) and the GTDB release 05-RS95 (Parks *et al*., 2018, 2020). MAGs assigned to the genus “UBA10452” in the order Nitrososphaerales (“terrestrial group I.1b”), phylum Thaumarchaeota (Rinke *et al*., 2021), were selected for downstream analyses **(Table 1, Suppl. Table S1)**. In addition, we analysed other eight UBA10452 MAGs available on GenBank and GTDB release 95 (Parks *et al*., 2018, 2020). These included six MAGs from permafrost soil in Canada (Chauhan *et al*., 2014; Parks *et al*., 2017), one MAG from polar desert soil in Antarctica (Ji *et al*., 2017), and one MAG from tundra soil in Finland (Pessi *et al*., 2022).

**Table 1.** List of metagenome-assembled genomes (MAGs) belonging to the UBA10452 lineage (*Candidatus* Nitrosopolaris).

| MAG | Isolation source | Accession | Ref. |
| --- | --- | --- | --- |
| COA_Bin_4_1 | Tundra soil, Rásttigáisá, Norway | GCA_933227015.1 | [1] |
| S89_Bin_2 | Tundra soil, Rásttigáisá, Norway | GCA_933227005.1 | [1] |
| S100_Bin_4 | Tundra soil, Rásttigáisá, Norway | GCA_933226995.1 | [1] |
| S1130_Bin_3 | Tundra soil, Rásttigáisá, Norway | GCA_933226985.1 | [1] |
| KWL-0179 | Tundra soil, Kilpisjärvi, Finland | GCA_936417005.1 | [2] |
| UBA272 | Permafrost soil, Nunavut, Canada | GCA_002504425.1 | [3] |
| UBA273 | Permafrost soil, Nunavut, Canada | GCA_002501935.1 | [3] |
| UBA347 | Permafrost soil (active layer), Nunavut, Canada | GCA_002495965.1 | [3] |
| UBA348 | Permafrost soil (active layer), Nunavut, Canada | GCA_002501855.1 | [3] |
| UBA466 | Permafrost soil (active layer), Nunavut, Canada | GCA_002498345.1 | [3] |
| UBA536 | Permafrost soil (active layer), Nunavut, Canada | GCA_002496625.1 | [3] |
| RRmetagenome_bin19 | Polar desert soil, Wilkes Land, Antarctica | GCA_003176995.1 | [4] |

1. This study.
2. Pessi et al. (2022, pre-print).
3. Parks et al. (2017), based on data originally published by Chauhan et al. (2014).
4. Ji et al. (2017).

### Genome annotation

We used anvi’o v7.0 (Eren *et al*., 2015) to predict gene calls with Prodigal v2.6.3 (Hyatt *et al*., 2010), identify ribosomal genes and a set of 76 archaeal single-copy genes with HMMER v.3.3 (Eddy, 2011), and compute genome completion and redundancy levels based on the presence of the 76 single-copy genes. We also employed anvi’o v7.0 to annotate the gene calls against the KOfam (Aramaki *et al*., 2020) and Pfam (Mistry *et al*., 2021) databases with HMMER v.3.3 (Eddy, 2011) and the COG database (Galperin *et al*., 2021) with DIAMOND v0.9.14 (Buchfink *et al*., 2015). Additionally, we used BLASTP v2.10.1 (Camacho *et al*., 2009) to annotate the gene calls against the arCOG database (Makarova *et al*., 2015). Matches with scores below the pre-computed family-specific thresholds (KOfam and Pfam), e-value > 10^−6^ (COG), or identity < 35% and coverage < 75% (arCOG) were discarded and, in case of multiple matches, the one with the lowest e-value was kept.

We used BLASTP v2.10.1 to compare the amino acid sequences of genes identified as *amoA, amoB*, or *amoC* to the RefSeq (O’Leary *et al*., 2016) and Swiss-Prot (The UniProt Consortium, 2019) databases, and BLASTN v2.10.1 (Camacho *et al*., 2009) to compare *amoA* genes against the nt database and the curated *amoA* database of Alves *et al*. (2018). Functional enrichment analyses were carried out using anvi’o v7.0 (Eren *et al*., 2015) according to Shaiber *et al*. (2020). In brief, the occurrence of arCOG functions across genomes was summarised and logistic regression was then used to identify functions associated with a particular genus or genera. For this, we considered only the three most complete *Nitrososphaera, Ca*. Nitrosocosmicus, and *Ca*. Nitrosodeserticola genomes plus the representative genome of each *Ca*. Nitrosopolaris species.

### Phylogenomic and phylogenetic analyses

For phylogenomic analysis, we used a set of 59 archaeal single-copy genes that were present in at least 80% of the genomes. In addition to the 12 UBA10452 MAGs, we retrieved from GenBank other 33 genomes belonging to the family Nitrososphaeraceae and the genome of *Nitrosopumilus maritimus* SCM1 to be used as an outgroup. We used anvi’o v7.0 (Eren *et al*., 2015) to recover the predicted amino acid sequence for each of the 59 genes, align them individually with MUSCLE v3.8.1551 (Edgar, 2004), and generate a concatenated alignment. We then computed a maximum likelihood tree with IQ-TREE v2.1.4 employing the automatic model selection and 1000 bootstraps (Nguyen *et al*., 2015). Pairwise average nucleotide identity (ANI) values were computed with pyani v0.2.10 (Pritchard *et al*., 2016) and amino acid identity (AAI) values with the AAI-Matrix tool (http://enve-omics.ce.gatech.edu/g-matrix).

Phylogenetic analysis of the *amoA* and 16S rRNA genes were done as described for the phylogenomic analysis (i.e., alignment with MUSCLE and tree building with IQ-TREE). Genes annotated as multicopper oxidase (PF07731, PF07732, COG2132, or arCOG03914) or nitrite reductase were aligned with MAFFT v7.490 (Katoh and Standley, 2013) alongside the sequences reported by Kerou *et al*. (2016), and a maximum likelihood tree was computed with IQ-TREE v2.1.4 (Nguyen *et al*., 2015) as described above.

### Abundance and geographic distribution

We employed read recruitment to compute the relative abundance of the UBA10452 lineage across the metagenomics datasets from which the MAGs were originally recovered. These datasets consisted of 10 Illumina NextSeq metagenomes from tundra soils in Rásttigáisá, Norway (this study); 69 Illumina NextSeq/NovaSeq metagenomes from tundra soils in Kilpisjärvi, Finland (Pessi *et al*., 2022, pre-print); 13 Illumina HiSeq metagenomes from permafrost soils in Nunavut, Canada (Chauhan *et al*., 2014; Stackhouse *et al*., 2015); and three Illumina HiSeq metagenomes from polar desert soils in Wilkes Land, Antarctica (Ji *et al*., 2017). We used fasterq-dump v2.10.8 (https://github.com/ncbi/sra-tools) to retrieve the raw metagenomic data from the Sequence Read Archive (SRA). We then used CoverM v0.6.1 (https://github.com/wwood/CoverM) to map the reads to the MAGs with minimap v2.17 (Li, 2016) and to compute relative abundances based on the proportion of reads recruited by the MAGs. In addition, we used IMNGS (Lagkouvardos *et al*., 2016) to further investigate the geographic distribution of the UBA10452 lineage. For this, we used the 16S rRNA gene sequence of the MAG RRmetagenome_bin19 as query to screen 422,877 amplicon sequencing datasets in SRA with UBLAST (Edgar, 2010). We considered only matches with ≥ 99.0% similarity.

## Results

### Genomic characteristics of the UBA10452 lineage

We applied a genome-resolved metagenomics approach to data obtained from tundra soils in Rásttigáisá, Norway, and recovered four MAGs assigned to the genus “UBA10452”, an uncultured lineage in the order Nitrososphaerales (“terrestrial group I.1b”), phylum Thaumarchaeota (Rinke *et al*., 2021). The UBA10452 lineage is currently represented by eight MAGs in GTDB and GenBank in addition to the four MAGs obtained in the present study **(Table 1, Fig. 1a)**. Genome completion and redundancy estimated with anvi’o v7.0 (Eren *et al*., 2015) based on the presence of 76 single-copy genes range from 50.0–90.8% and 2.6–9.2%, respectively **(Fig. 1b, Suppl. Table S1)**. The MAG RRmetagenome_bin19, with 90.8% completion, 3.9% redundancy, and a nearly complete (1462 bp) 16S rRNA gene, is a high-quality MAG according to the MIMAG standard (Bowers *et al*., 2017). The remaining 11 MAGs are of medium quality (≥ 50% complete, < 10% redundant), four of which also include the 16S rRNA gene. The genome size of UBA10452 MAGs ranges from 0.8 Mb (MAG S1130_Bin_3, 60.5% complete) to 4.0 Mb (MAG RRmetagenome_bin19, 90.8% complete). G+C content ranges from 38.1 to 41.5%.

**Figure 1.**
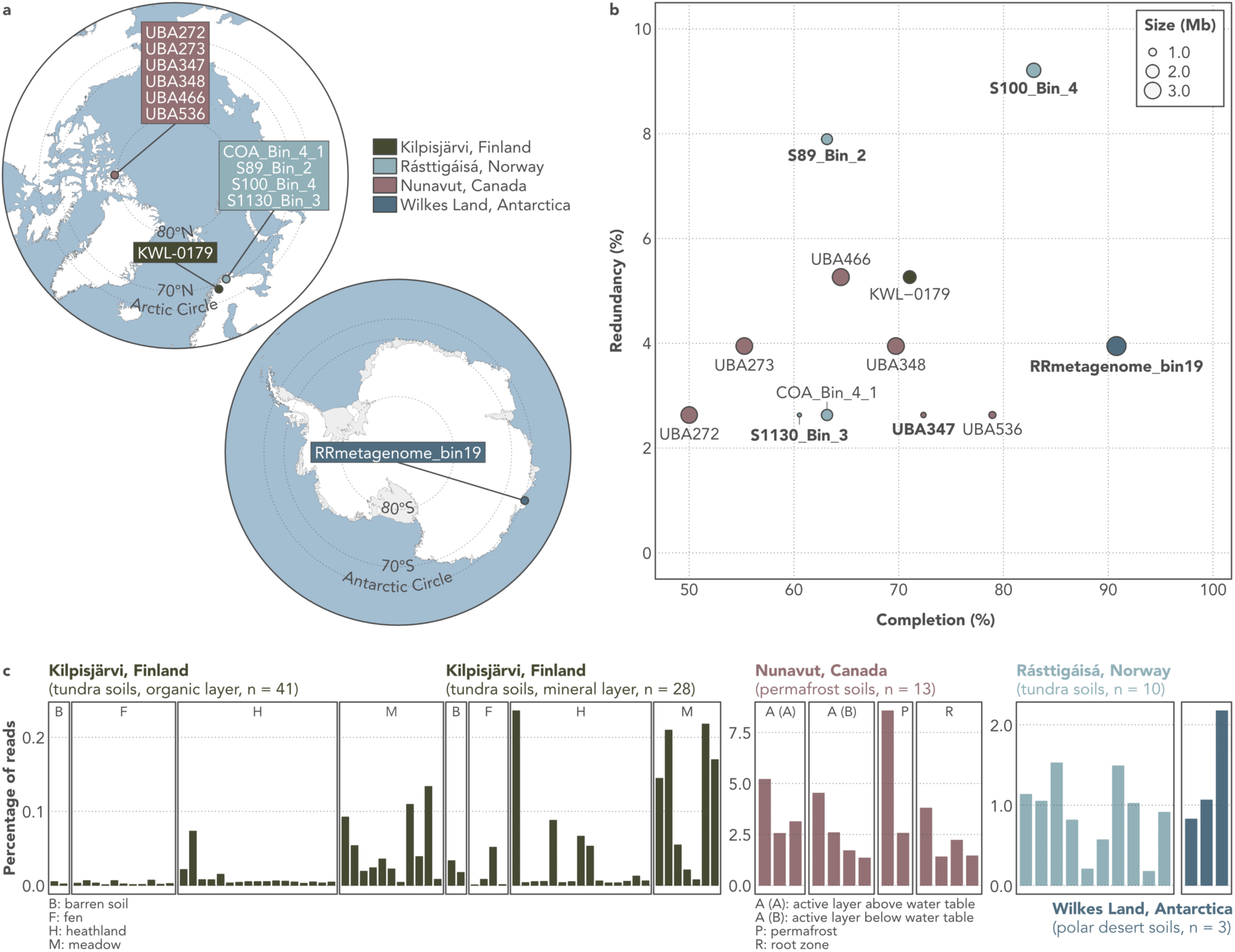
Geographic origin, assembly statistics, and abundance of metagenome-assembled genomes (MAGs) assigned to the UBA10452 lineage (*Candidatus* Nitrosopolaris). **a)** Maps of the Arctic and Antarctic regions showing the geographic origin of the 12 UBA10452 MAGs. **b)** Genome completion, redundancy, and size of the UBA10452 MAGs. Completion and redundancy levels were computed based on the presence of 76 single-copy genes. MAGs in bold include the 16S rRNA gene. **C)** Proportion of metagenomic reads recruited by the UBA10452 MAGs across the four datasets from which they were originally recovered.

### UBA10452 has a predominantly polar distribution

All 12 UBA10452 MAGs were obtained from tundra, permafrost, and polar desert soils **(Table 1, Fig. 1a)**. To gain insights into the ecology of the UBA10452 lineage, we used read recruitment to quantify the abundance of UBA10452 MAGs in the metagenomic datasets from which they were assembled. UBA10452 MAGs were most abundant in the dataset of permafrost from nutrient-poor (C: 1.0%, N: 0.1%) mineral cryosoils in Nunavut, Canadian Arctic, where they recruited up to 8.6% of the reads in each sample **(Fig. 1c)**. On the other hand, UBA10452 MAGs were least abundant in the more nutrient-rich (C: 7.3%, N: 0.3%) tundra soils from Kilpisjärvi, Finland, where they were detected particularly in samples taken from the mineral layer of heathland and meadow soils.

In order to investigate further the geographic distribution of the UBA10452 lineage, we used IMNGS (Lagkouvardos *et al*., 2016) to screen 422,877 16S rRNA gene amplicon sequencing datasets in SRA. Sequences matching the 16S rRNA gene of UBA10452 MAGs (≥ 99.0% similarity) were found across 1281 datasets, mostly consisting of soil (n = 750), freshwater (n = 104), and rhizosphere samples (n = 100). Matched reads accounted for 6.0% of the total number of reads in these datasets (8.9 out of 149.1 million sequences). Of these, the overwhelming majority (8.7 million reads, 97.9%) come from Antarctic soil datasets, particularly from 149 sites in the vicinity of Davis Station, Princess Elizabeth Land (Bissett *et al*., 2016) **(Suppl. Fig. S1a)**. The proportion of reads matching the UBA10452 lineage was above 50% of the archaeal 16S rRNA gene sequences in 70 of these sites and reached values as high as 88.8% **(Suppl. Fig. S1b)**.

### UBA10452 is a distinct lineage in the family Nitrososphaeraceae

Phylogenomic analysis based on 59 single2-copy genes placed the UBA10452 MAGs as a distinct lineage outside *Nitrososphaera, Ca*. Nitrosocosmicus, and *Ca*. Nitrosodeserticola, the three described genera in the family Nitrososphaeraceae (Stieglmeier *et al*., 2014; Lehtovirta-Morley *et al*., 2016; Hwang *et al*., 2021) **(Fig. 2a; Suppl. Fig. S2)**. Separation of UBA10452 is also supported by AAI and 16S rRNA gene analyses. UBA10452 MAGs share 59.1% ± 1.9, 53.0% ± 1.1, and 53.8% ± 0.9 AAI with *Nitrososphaera, Ca*. Nitrosocosmicus, and *Ca*. Nitrosodeserticola, respectively **(Fig. 2b)**, all of which are below the 65% AAI threshold commonly used to delineate microbial genera (Konstantinidis *et al*., 2017). At the 16S rRNA gene level, UBA10452 MAGs are 94.8% ± 1.2 and 95.4% ± 0.2 similar to *Nitrososphaera* and *Ca*. Nitrosocosmicus, respectively **(Suppl. Fig. S3)**. These values are in the limit of the 95% threshold for genus delineation proposed by Rosselló-Móra and Amann (2015), but are well below the median 16S rRNA gene similarity observed between related genera across different microbial phyla (96.4%; Yarza *et al*., 2014). Comparison with *Ca*. Nitrosodeserticola was not possible due to the lack of a 16S rRNA gene sequence from this genus. Given that UBA10452 represents a clear, distinct lineage in the family Nitrososphaeraceae, we consider that UBA10452 should be recognized as a *Candidatus* genus and propose the name *Ca*. Nitrosopolaris.

**Figure 2.**
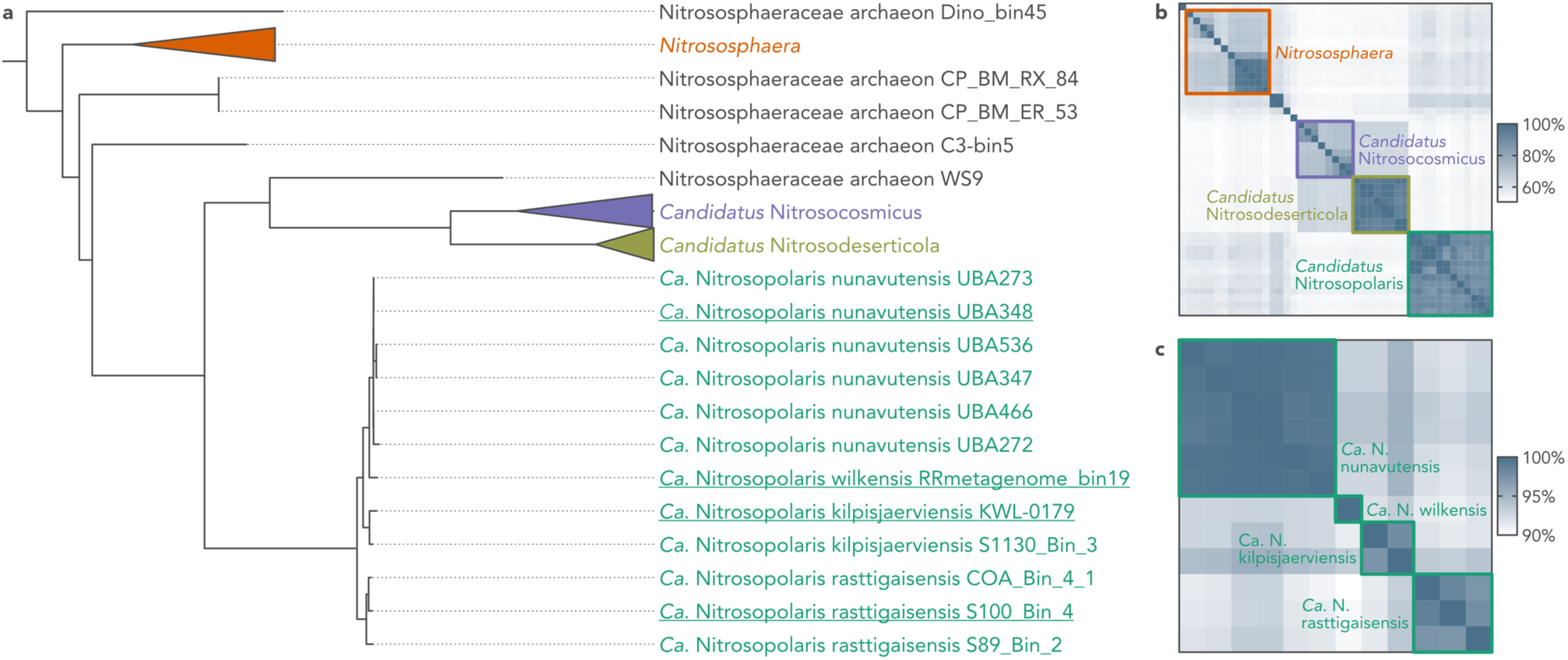
Phylogenomic analysis of the UBA10452 lineage (*Candidatus* Nitrosopolaris). **a)** Maximum likelihood tree based on 59 single-copy genes from 12 metagenome-assembled genomes (MAGs) assigned to the UBA10452 lineage and 33 other Nitrososphaeraceae genomes available on GenBank. The tree was rooted with *Nitrosopumilus maritimus* SCM1 (not shown). Representatives for the four proposed species are indicated in underscore. An uncollapsed and bootstrapped version of the tree can be found in **Suppl. Fig. S2. b)** Pairwise average amino acid identity (AAI) between Nitrososphaeraceae genomes and **c)** average nucleotide identity (ANI) between UBA10452 MAGs. The boxes encompass the four described Nitrososphaeraceae genera (AAI threshold of 65%; panel b) and the four proposed species of *Candidatus* Nitrosopolaris (ANI threshold of 95–96%; panel c). Rows and columns are ordered from top to bottom and left to right, respectively, according to the top-bottom order of leaves in panel a.

Pairwise ANI values between *Ca*. Nitrosopolaris MAGs range from 90.9 to 99.9% **(Fig. 2c)**. Based on either a 95% (Konstantinidis *et al*., 2017) or 96% ANI threshold (Ciufo *et al*., 2018), the 12 *Ca*. Nitrosopolaris MAGs can be separated into four distinct species **(Fig. 2a; Suppl. Fig. S2)**. Two of these, one comprising the six Canadian MAGs (Chauhan *et al*., 2014; Parks *et al*., 2017) and the other consisting solely of the Antarctic MAG (Ji *et al*., 2017), correspond to the two existing species in GTDB release 95 (“UBA10452 sp002501855” and “UBA10452 sp003176995”, respectively). Here we suggest renaming these species as *Ca*. Nitrosopolaris nunavutensis and *Ca*. Nitrosopolaris wilkensis, respectively, according to the geographic origin of the MAGs. The Finnish MAG (Pessi *et al*., 2022, pre-print) plus one of the Norwegian MAGs obtained in the present study (S1130_Bin_3) represent a novel species, for which we suggest the name *Ca*. Nitrosopolaris kilpisjaerviensis. Finally, the three remaining MAGs obtained in the present study (COA_Bin_4_1, S89_Bin_2, and S100_Bin_4) correspond to another novel species, which we propose to be named as *Ca*. Nitrosopolaris rasttigaisensis. However, the separation of *Ca*. Nitrosopolaris into four species is not supported by the analysis of the 16S rRNA gene **(Suppl. Fig. S3)**. The pairwise similarity between 16S rRNA gene sequences across the four ANI clusters range from 99.5 to 99.9%, which is above the 98.7–99.0% threshold commonly used for species delineation (Stackebrandt and Ebers, 2006; Kim *et al*., 2014).

### *Ca*. Nitrosopolaris harbours genes for ammonia oxidation, CO_2_ fixation, and carbohydrate metabolism and transport

Annotation of protein-coding genes revealed that *Ca*. Nitrosopolaris harbours the *amoA, amoB, amoC*, and *amoX* genes encoding the enzyme ammonia monooxygenase (Amo) which catalyses the oxidation of ammonia to hydroxylamine **(Fig. 3a, Suppl. Fig. S4, Suppl. Table S2)**. As for other AOA, homologues of the *hao* gene were not found in the *Ca*. Nitrosopolaris MAGs. In AOB, this gene encodes the enzyme hydroxylamine dehydrogenase (Hao) which takes part in the oxidation of hydroxylamine to nitrite, a mechanism that remains unknown in AOA (Lehtovirta-Morley, 2018). A proposed mechanism of hydroxylamine oxidation in AOA is via a copper-containing nitrite reductase (NirK) encoded by the *nirK* gene, which has been detected in most *Ca*. Nitrosopolaris MAGs as well as other related multicopper oxidases **(Suppl. Fig. S4)**. *Ca*. Nitrosopolaris also encodes an ammonium transporter of the Amt family involved in the uptake of extracellular ammonium. Moreover, *Ca*. Nitrosopolaris harbours urease (*ureABC*) and urea transporter (*utp*) genes, indicating the ability to generate ammonia from urea. In contrast to *Nitrososphaera gargensis* (Spang *et al*., 2012), we did not detect the *cynS* gene encoding the enzyme cyanate hydratase involved in the production of ammonia from cyanate.

**Figure 3.**
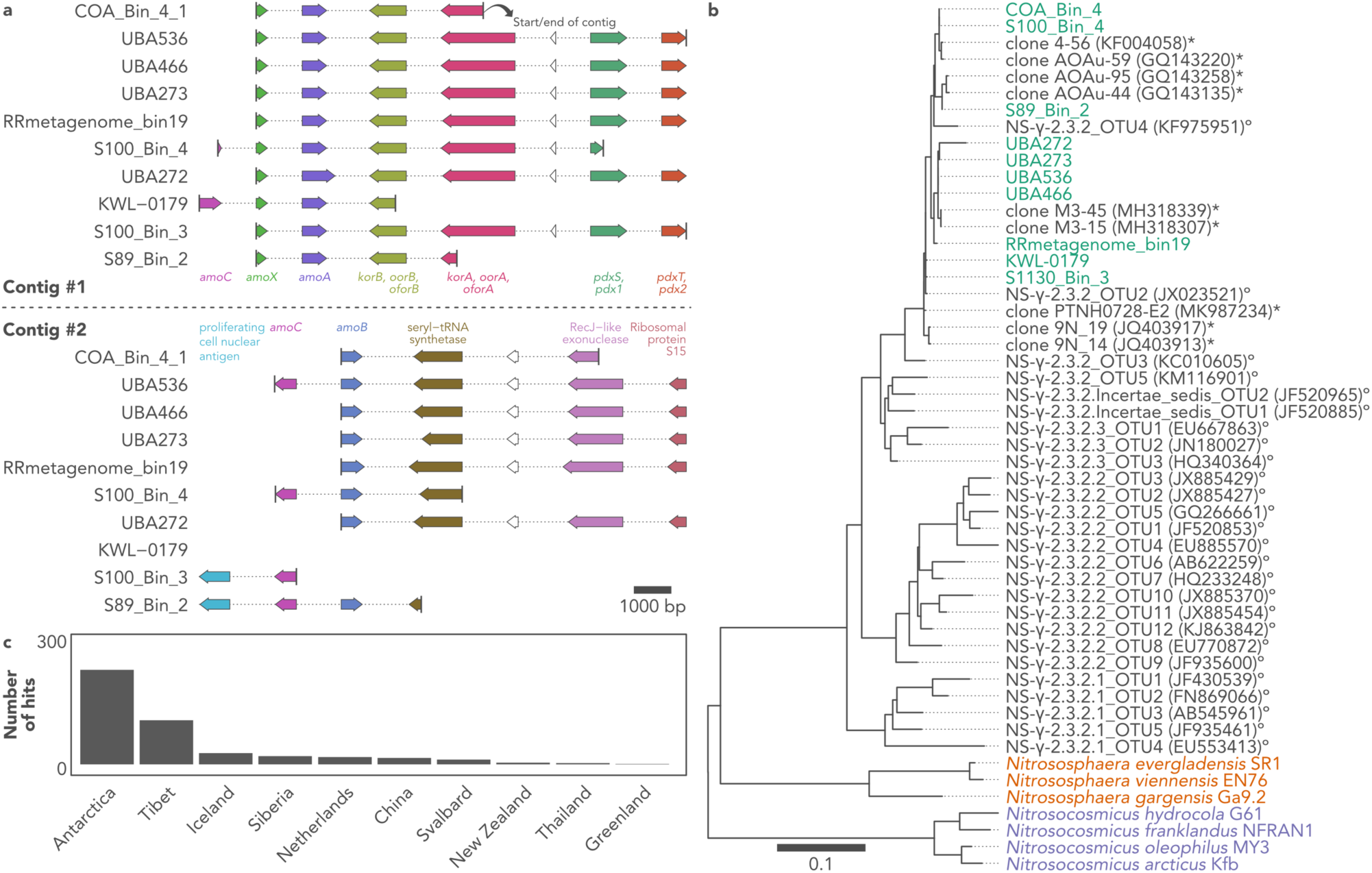
The ammonia monooxygenase (*amo*) genes of UBA10452 (*Candidatus* Nitrosopolaris). **a)** Representation of two contigs containing *amo* genes in metagenome-assembled genomes (MAGs) assigned to the UBA10452 lineage. Two MAGs which do not contain the *amoA* gene are omitted (UBA347 and UBA348). **b)** Maximum likelihood tree of the *amoA* sequence of UBA10452 MAGs and related sequences from GenBank (asterisks) and Alves *et al*. (2018) (circles). **c)** Geographic origin of sequences from GenBank with ≥ 96.0% nucleotide similarity to the *amoA* sequence of UBA10452 MAGs.

The *amo* genes in all *Ca*. Nitrosopolaris MAGs are distributed across two separate contigs **(Fig. 3a)**. One of the contigs contains the *amoC, amoX*, and *amoA* genes; however, the *amoC* gene is truncated and found in only two MAGs. In some MAGs, the other contig contains a second, full-length copy of the *amoC* gene followed by *amoB*. Not all MAGs contain all *amoABCX* genes. However, considering that the MAGs present varying levels of completion **(Fig. 1b, Suppl. Table S1)** and since the localization of the genes corresponds to start or end of contigs **(Fig. 3a)**, it is likely that missing genes are an artifact of truncated assemblies rather than due to gene loss. The *amoA* gene of *Ca*. Nitrosopolaris has a length of 651 bp and belongs to the NS-γ-2.3.2 cluster of Alves *et al*. (2018) **(Fig. 3b)**. One exception is the MAG UBA272 which contains a longer *amoA* gene (873 bp) with a long insert of ambiguous base calls, most likely an artifact from assembling and/or scaffolding. Sequences belonging to the NS-γ-2.3.2 cluster are found majorly in acidic soils (Alves *et al*., 2018). Moreover, analysis of sequences from GenBank showed that the *amoA* gene of *Ca*. Nitrosopolaris is related (≥ 96% nucleotide similarity) to sequences recovered mostly from cold environments, i.e., the Artic, Antarctica, and alpine regions such as the Tibetan Plateau **(Fig. 3c)**. Among these, the *amoA* sequences of *Ca*.

Nitrosopolaris MAGs are most closely related (98.9–99.7% nucleotide similarity) to uncultured sequences from Antarctic soil (MH318339 and MH318307), grassland soil in Iceland (JQ403917 and JQ403913), and the Tibetan Plateau (GQ143258, GQ143220, GQ143135, KF004058, and MK987234) (Daebeler *et al*., 2012; Xie *et al*., 2014; Wang *et al*., 2019; Zhang *et al*., 2019) **(Fig. 3b)**.

Similarly to other AOA, *Ca*. Nitrosopolaris harbours genes for the hydroxypropionate-hydroxybutyrate pathway of CO_2_ fixation, complexes I–V of the electron transfer chain, the citric acid cycle, and gluconeogenesis **(Suppl. Fig. S4, Suppl. Table S2)**. Like other AOA, the gene content of *Ca*. Nitrosopolaris indicates a potential for mixotrophic metabolism, with multiple copies of genes encoding proteins involved in carbohydrate metabolism and transport such as glucose/sorbosone dehydrogenases, permeases of the major facilitator superfamily (MFS), and pyruvate oxidases. In contrast to *Nitrososphaera*, we did not detect genes involved in the assembly of pili, flagellar apparatus (archaellum), and chemotaxis.

### *Ca*. Nitrosopolaris MAGs are enriched in genes involved in DNA replication and repair

To investigate possible mechanisms underlying the distribution of *Ca*. Nitrosopolaris, we carried out a functional enrichment analysis covering the four Nitrososphaeraceae genera. In total, the 13 MAGs used in the analysis encoded 3,999 different arCOG functions **(Fig. 4a)**. Of these, 948 functions were shared among all four genera and 368 were unique to *Ca*. Nitrosopolaris. Of the arCOG functions shared by all four genera, most belonged to the arCOG classes translation, ribosomal structure, and biogenesis (n = 114), function unknown (n = 88), and amino acid transport and metabolism (n = 73). On the other hand, arCOG functions unique to *Ca*. Nitrosopolaris belonged mostly to the arCOG classes function unknown (n = 59), general function prediction only (n = 52), and inorganic ion transport and metabolism (n = 33). Among these arCOG functions are several types of hydrolases, lipoproteins, phospholipases, and ABC transporters including ones for iron, maltose, phosphate, amino acids, and nucleosides **(Suppl. Table S3)**.

**Figure 4.**
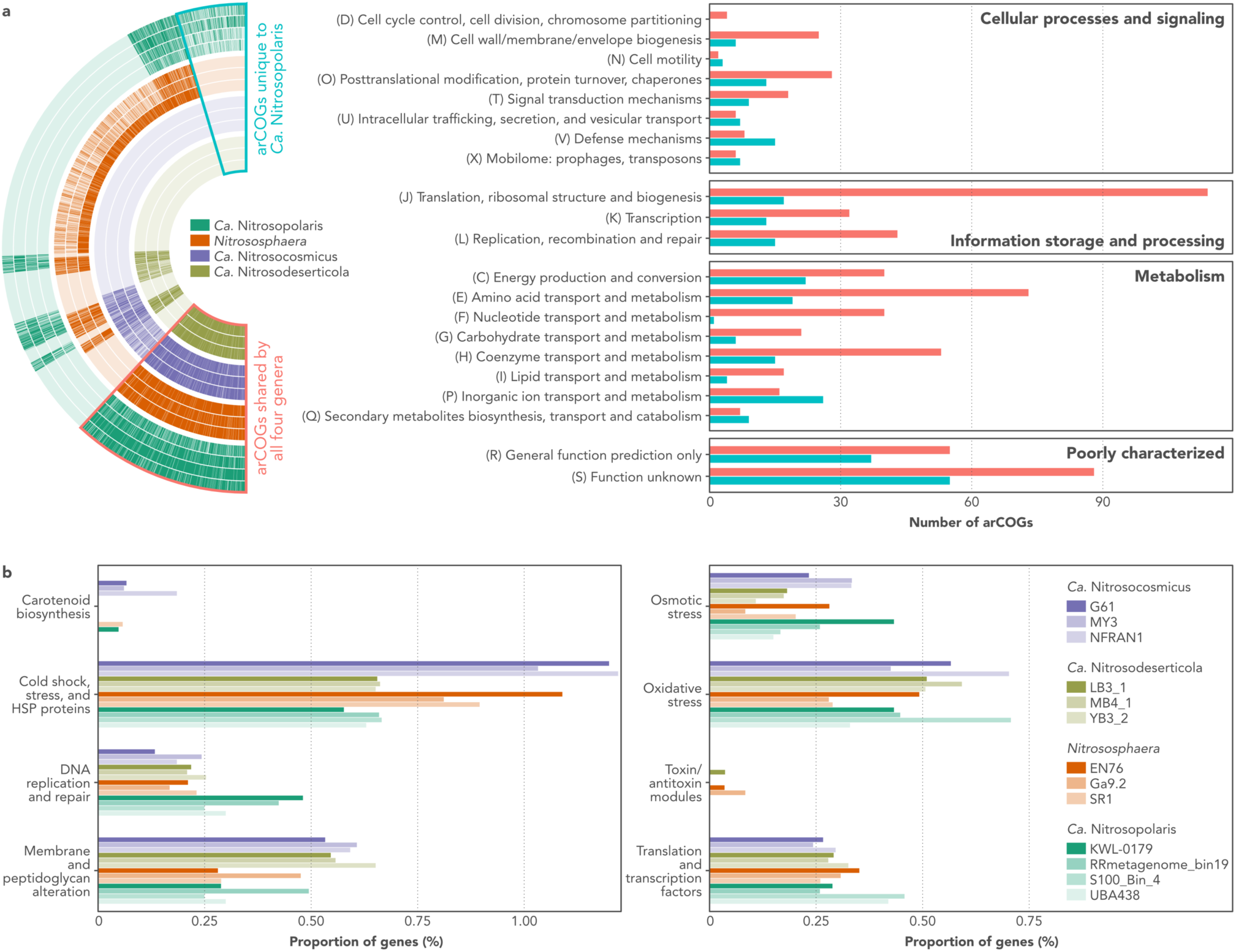
Comparative genomics of the UBA10452 lineage (*Candidatus* Nitrosopolaris) and other members of the family Nitrososphaeraceae. **a)** Functional enrichment analysis showing arCOG functions shared by all Nitrososphaeraceae genera and other functions unique to *Ca*. Nitrosopolaris. More detail on the arCOG functions unique to *Ca*. Nitrosopolaris can be found in **Suppl. Table S3. b)** Distribution of genes with known or predicted roles in cold adaptation and growth (Raymond-Bouchard *et al*., 2018). Number of genes is shown as a proportion of the total number of genes in each genome. More detail on the genes found in *Ca*. Nitrosopolaris can be found in **Suppl. Table S4**.

In addition to the genome-wide functional enrichment analysis, we also looked more specifically for genes with known or predicted roles in cold adaptation and growth (Raymond-Bouchard *et al*., 2018). When comparing the genomic repertoire of *Ca*. Nitrosopolaris to the other Nitrososphaeraceae genera, the former was found to harbour a higher number of genes involved in DNA replication and repair **(Fig. 4b)**. More specifically, *Ca*. Nitrosopolaris MAGs encode multiple copies of the enzymes RecA ATPases and RecA/RadA recombinases **(Suppl. Table S4)**. Surprisingly, genes related to cold shock response were less abundant in *Ca*. Nitrosopolaris compared to *Nitrososphaera* and *Ca*. Nitrosocosmicus **(Fig. 4b)**, although several copies of molecular chaperones (DnaK, GrpE, and IbpA) and universal stress proteins (UspA) were identified **(Suppl. Table S4)**. In addition to these, *Ca*. Nitrosopolaris also harbours several copies of other genes encoding proteins related to cold adaptation and growth **(Fig. 4b)**, including proteins involved in membrane and peptidoglycan alteration (glycosyltransferases), osmotic stress (sodium-hydrogen antiporters and sodium-proline symporters), oxidative stress (periredoxins and thioredoxin reductases), and translation/transcription (DNA/RNA helicases and transcription factors) **(Suppl. Table S4)**.

## Discussion

Genome-resolved metagenomics has revolutionized our knowledge of archaeal diversity by giving us access to the genome of uncultured microorganisms at an unprecedented rate (Tahon *et al*., 2021). In a recent metagenomic investigation of tundra soils in northern Finland (Pessi *et al*., 2022, pre-print), we have manually binned and curated a MAG belonging to the genus “UBA10452”, an uncultured and largely uncharacterized lineage in the order Nitrososphaerales (“terrestrial group I.1b”) of the phylum Thaumarchaeota (Rinke *et al*., 2021). Here, we binned four other UBA10452 MAGs from tundra soils in Rásttigáisá, Norway, and characterized the phylogeny, metabolic potential, and biogeography of this lineage. Our results indicate that the UBA10452 lineage consists of putative AOA with a geographic distribution mostly restricted to cold ecosystems, particularly the polar regions. We suggest the recognition of UBA10452 as a *Candidatus* genus, for which we propose the name *Ca*. Nitrosopolaris (*nitrosus*: Latin adjective meaning nitrous; *polaris*: Latin adjective meaning of or pertaining to the poles).

The findings from our polyphasic analysis consisting of phylogenomic, AAI, and 16S rRNA gene analyses support the placement of *Ca*. Nitrosopolaris outside *Nitrososphaera, Ca*. Nitrosocosmicus, and *Ca*. Nitrosodeserticola in the family Nitrososphaeraceae, as previously suggested (Rinke *et al*., 2021). Our results further indicate that the 12 *Ca*. Nitrosopolaris MAGs represent four different species based on a 95–96% ANI threshold (Konstantinidis *et al*., 2017; Ciufo *et al*., 2018). In addition to the two current species in GTDB release 95 (Parks *et al*., 2018, 2020), the inclusion of the four MAGs obtained in the present study resulted in the identification of two novel species. It is important to note that the separation of *Ca*. Nitrosopolaris into four species based on ANI values is not readily supported by the analysis of 16S rRNA gene sequences, which are ≥ 99.5% similar across the four species. Although a 98.7–99.0% threshold is commonly used (Stackebrandt and Ebers, 2006), species delineation based solely on the 16S rRNA gene can be problematic given that microorganisms belonging to different species can share identical 16S rRNA gene sequences (Kim *et al*., 2014; Schloss, 2021). One example of this is *Ca*. Nitrosocosmicus arcticus and *Ca*. Nitrosocosmicus oleophilus, two species of AOA which share an identical 16S rRNA gene sequence despite having divergent genomes with only 83.0% ANI (Alves *et al*., 2019). It thus appears reasonable to conclude that the 12 *Ca*. Nitrosopolaris MAGs indeed represent four different species as suggested by the ANI analysis. If cultured representatives become available in the future, phenotypic and ecophysiological characterization of these isolates could help resolve the taxonomy of *Ca*. Nitrosopolaris.

*Ca*. Nitrosopolaris harbours the complete set of *amoA* genes responsible for chemolithotrophic growth via ammonia oxidation (Lehtovirta-Morley, 2018). Although *in silico* analyses provide valuable predictions, metabolic capabilities inferred by genomic annotation need to be confirmed based on the analysis of isolated/enriched cultures or with the help of other indirect methods such as stable isotope probing (SIP) (Gadkari *et al*., 2020). Nevertheless, the putative ammonia oxidation capability of *Ca*. Nitrosopolaris is supported by its close phylogenetic relationship to *Nitrososphaera* and *Ca*. Nitrosocosmicus, two genera which have been demonstrated to grow by oxidizing ammonia (Stieglmeier *et al*., 2014; Lehtovirta-Morley *et al*., 2016). In addition, the presence of several genes involved in carbohydrate and amino acid transport and metabolism suggest that *Ca*. Nitrosopolaris, like other AOA, might be able to grow mixotrophically using organic compounds as alternative energy and/or C sources (Mussmann *et al*., 2011; Pester *et al*., 2011).

In addition to the geographical origin of the MAGs, large-scale screening of 16S rRNA gene and *amoA* sequences from SRA and GenBank indicate that *Ca*. Nitrosopolaris is restricted to soils in the cold biosphere. The soils from which the *Ca*. Nitrosopolaris MAGs have been recovered are typical of polar and alpine environments, being characterized by low pH (4.8–5.1), carbon (C) (1.0–7.3%), and N (0.1–0.3%) content (Stackhouse *et al*., 2015; Ji *et al*., 2017; Pessi *et al*., 2022, pre-print). Furthermore, the abundance profile of *Ca*. Nitrosopolaris observed in this study, which was characterized by a higher abundance in mineral cryosoil permafrost and polar desert soils compared to vegetated tundra soils, indicates that *Ca*. Nitrosopolaris is particularly adapted to the highly oligotrophic conditions found in some of the most extreme environments in the cryosphere. The discovery of *Ca*. Nitrosopolaris complements the list of microbial taxa that appear to be adapted to life in cold environments, such as the mat-forming cyanobacteria *Phormidesmis priestleyi* (Komárek *et al*., 2009) and *Shackeltoniella antarctica* (Strunecky *et al*., 2020) and the sea-ice bacteria *Polaribacter* and *Psychrobacter* (Bowman, 2013).

Investigation of the genome of *Ca*. Nitrosopolaris provided insights on possible adaptations to cold and oligotrophic environments. For instance, *Ca*. Nitrosopolaris harbour multiple copies of several genes that have been implicated in tolerance to cold, such as genes encoding proteins involved in DNA replication and repair, molecular chaperones, DNA/RNA helicases, and universal stress proteins (Raymond-Bouchard *et al*., 2018). Interestingly, *Ca*. Nitrosopolaris appears to be enriched in copies of the RecA enzyme compared to other members of the Nitrososphaeraceae. RecA plays a key role in DNA repair, which is an important mechanism for survival in polar environments where DNA is frequently damaged due to freezing and UV radiation (Cavicchioli, 2006). In addition to the possible adaptive mechanisms of *Ca*. Nitrosopolaris, it has been suggested that the environmental characteristics of polar soils favour AOA in general (Alves *et al*., 2013; Siljanen *et al*., 2019). The ecological success of AOA in oligotrophic and acidic soils has been traditionally linked to the higher affinity of their ammonia oxidation machinery compared to their bacterial counterparts (Martens-Habbena *et al*., 2009; Kerou *et al*., 2016), although a recent study has shown that high affinity for ammonia is not common to all AOA (Jung *et al*., 2022). Furthermore, the hydroxypropionate-hydroxybutylate pathway of CO_2_ fixation encoded by *Ca*. Nitrosopolaris and other AOA appears to be more energy efficient than the Calvin cycle employed by AOB (Könneke *et al*., 2014). However, these traits are shared between *Ca*. Nitrosopolaris and other AOA and thus do not readily explain the apparent ecological success of *Ca*. Nitrosopolaris in cold environments. Indeed, mechanisms of cold adaptation are evolutionary and functionally complex and involve many features that cannot be observed by metagenomics alone (e.g., gene regulation and membrane modifications) (Cavicchioli, 2006). Structural, transcriptomics, and proteomics analysis of cultured isolates could help shed further light on possible adaptations to cold in *Ca*. Nitrosopolaris.

In addition to possible mechanisms of adaptation to polar environments, we hypothesize that the distribution of *Ca*. Nitrosopolaris could be, to some extent, related to historical factors. Interestingly, the four proposed *Ca*. Nitrosopolaris species form coherent biogeographical clusters: *Ca*. N. nunavutensis, comprising MAGs obtained from permafrost soils in Nunavut, Canada; *Ca*. N. wilkensis, corresponding to one MAG from polar desert soils in Wilkes Land, Antarctica; and *Ca*. N. kilpisjaerviensis and *Ca*. N. rasttigaisensis, comprising MAGs obtained from mineral tundra soils in two relatively close regions in northern Fennoscandia (Kilpisjärvi and Rásttigáisá, respectively). A recent molecular dating study has suggested that the origin of the AOA clade group I.1b (order Nitrososphaerales) coincides with severe glaciation events that happened during the Neoproterozoic (Yang *et al*., 2021). If these estimates are accurate, it would imply that *Ca*. Nitrosopolaris and all other lineages in group I.1b share a common ancestor that appeared when the global climate was characterized by sub-zero temperatures, having likely evolved at glacial refugia such as nunataks or regions with geothermal activity.

Due to low temperatures throughout the year, polar soils store a large amount of organic matter and have thus served as important carbon sinks. At present, polar soils are considered minor yet significant sources of N_2_O (Voigt *et al*., 2020) but, if warming trends continue at the levels observed currently, polar soils might become major contributors to the global N_2_O budget. For instance, the AOA *Ca*. Nitrosocosmicus arcticus isolated from Arctic soil has an ammonia oxidation optimum at temperatures well above those found *in situ* (Alves *et al*., 2019). Given that both the direct and indirect roles of AOA in the cycling of N_2_O in polar soils remain largely undetermined, a better understanding of polar microbial communities is paramount to model current and future N_2_O fluxes from this biome.

## Acknowledgments

This study was funded by the Academy of Finland (project 335354) and the University of Helsinki. We would like to acknowledge the CSC – IT Centre for Science, Finland, for providing computing resources; Miska Luoto and the BioGeoClimate Modelling Lab at the University of Helsinki for providing the Rásttigáisá samples; Eeva Eronen-Rasimus, Laura Cappelatti, and Benoit Durieu for helpful suggestions and discussions; and Jouko Rikkinen and Heino Vänskä for nomenclatural advice.

## Data availability

Genomic assemblies can be found in GenBank/ENA under the accession numbers listed in **Table 1**. MAGs generated in this study have been submitted to ENA (BioProject PRJEB49283). All the code used can be found in https://github.com/ArcticMicrobialEcology/candidatus-nitrosopolaris.

## Author contributions

ISP, AR, and JH designed the study. ISP performed the analyses and wrote the manuscript. AR and JH revised the manuscript.

## SSupplementary Information

**Suppl. Table S1 (separate** .**xlsx file)**. Additional information on metagenome-assembled genomes (MAGs) belonging to the UBA10452 lineage (*Candidatus* Nitrosopolaris).

**Suppl. Table S2 (separate** .**xlsx file)**. Information about selected genes used for the reconstruction of the metabolic potential of the UBA10452 lineage (*Candidatus* Nitrosopolaris).

**Suppl. Table S3 (separate** .**xlsx file)**. arCOG functions enriched in metagenome-assembled genomes (MAGs) belonging to the UBA10452 lineage (*Candidatus* Nitrosopolaris).

**Suppl. Table S4 (separate** .**xlsx file)**. Genes with known or predicted roles in cold adaptation and growth found in metagenome-assembled genomes (MAGs) belonging to the UBA10452 lineage (*Candidatus* Nitrosopolaris).

**Suppl. Figure S1.**
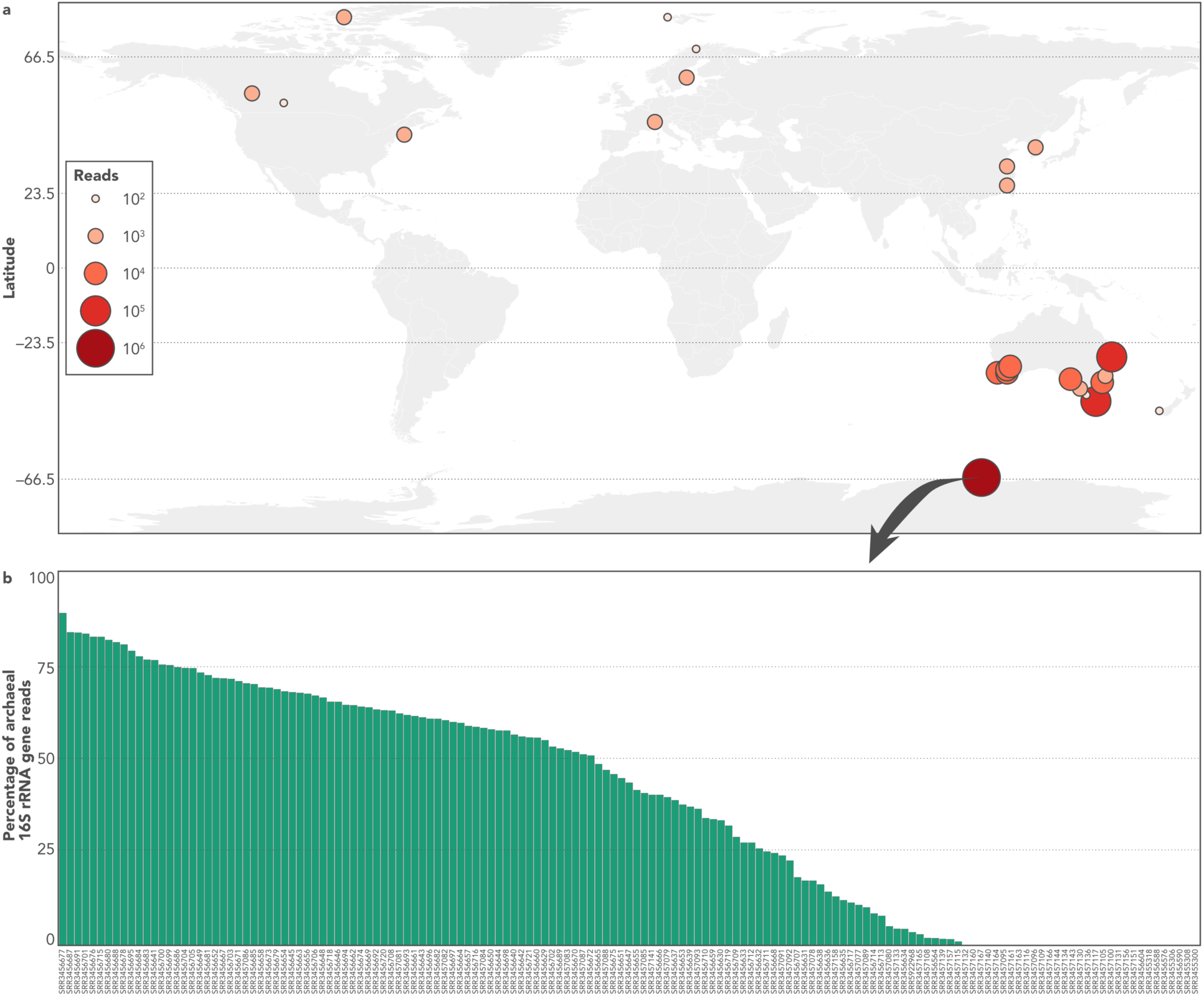
Geographic distribution of the UBA10452 lineage (*Candidatus* Nitrosopolaris). **a)** Distribution of *Ca*. Nitrosopolaris based on the screening of 422,877 16S rRNA gene amplicon sequencing datasets in the Sequence Read Archive (SRA). Datasets with few matches (< 0.1% or < 100 reads) are not shown. **b)** Abundance of *Ca*. Nitrosopolaris across 149 16S rRNA gene amplicon sequencing datasets from soils in the vicinity of Davis Station, Princess Elizabeth Land, Antarctica (BioProject PRJNA317932). Relative abundances were computed as the proportion of reads matching the sequence of *Ca*. Nitrosopolaris in each sample. Abundances represent the percentage of *Ca*. Nitrosopolaris reads relative to archaeal 16S rRNA gene reads obtained with archaea-specific primers.

**Suppl. Figure S2.**
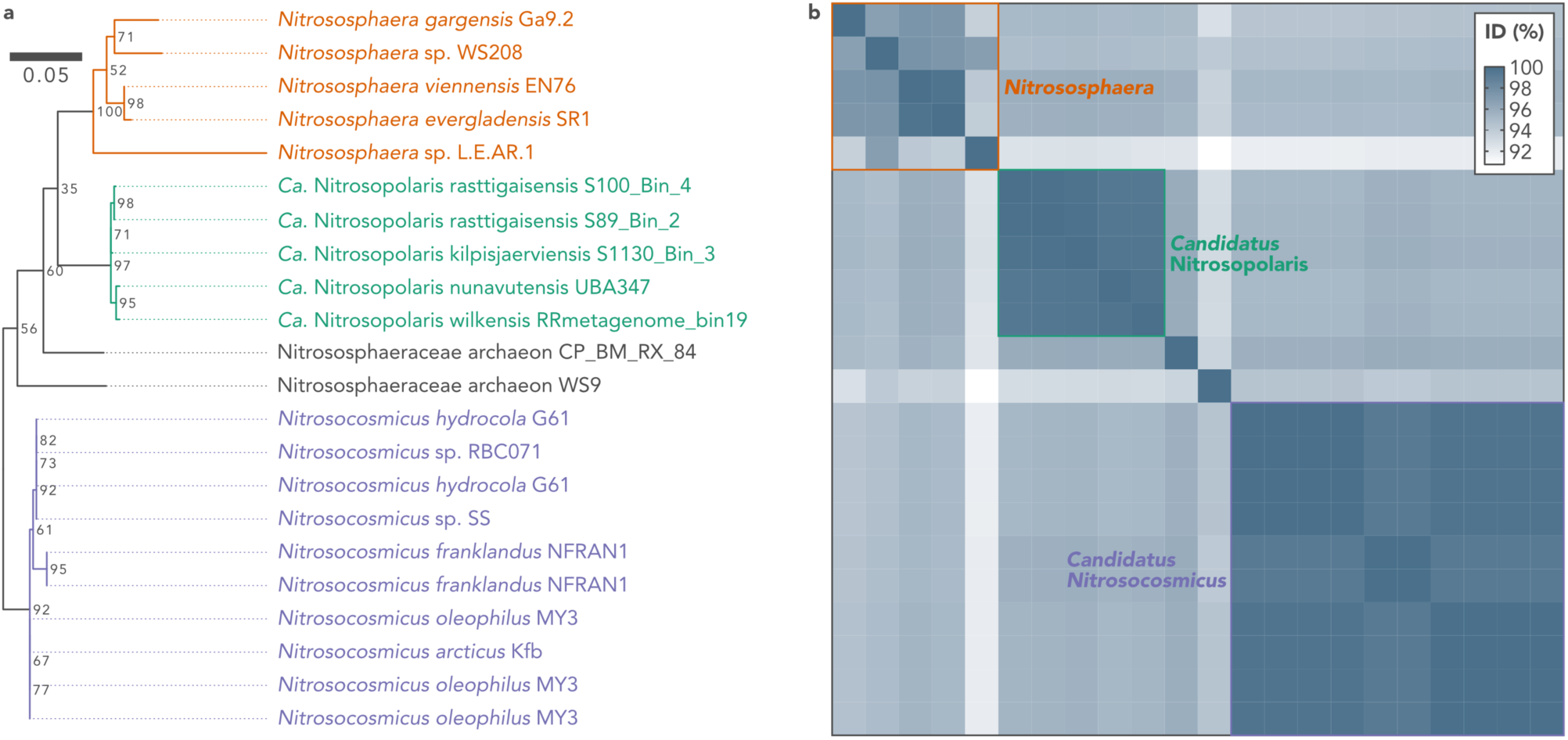
The 16S rRNA gene of UBA10452 (*Candidatus* Nitrosopolaris). **a)** Phylogenetic analysis of the 16S rRNA gene sequence of five metagenome-assembled genomes (MAGs) assigned to the UBA10452 lineage and other Nitrososphaeraceae genomes available on GenBank. Maximum likelihood tree rooted with *Nitrosopumilus maritimus* SCM1 (not shown). Bootstrap values for node support are indicated. **b)** Pairwise similarity between 16S rRNA gene sequences. Note that some genomes contain multiple copies of the 16S rRNA gene.

**Suppl. Figure S3.**
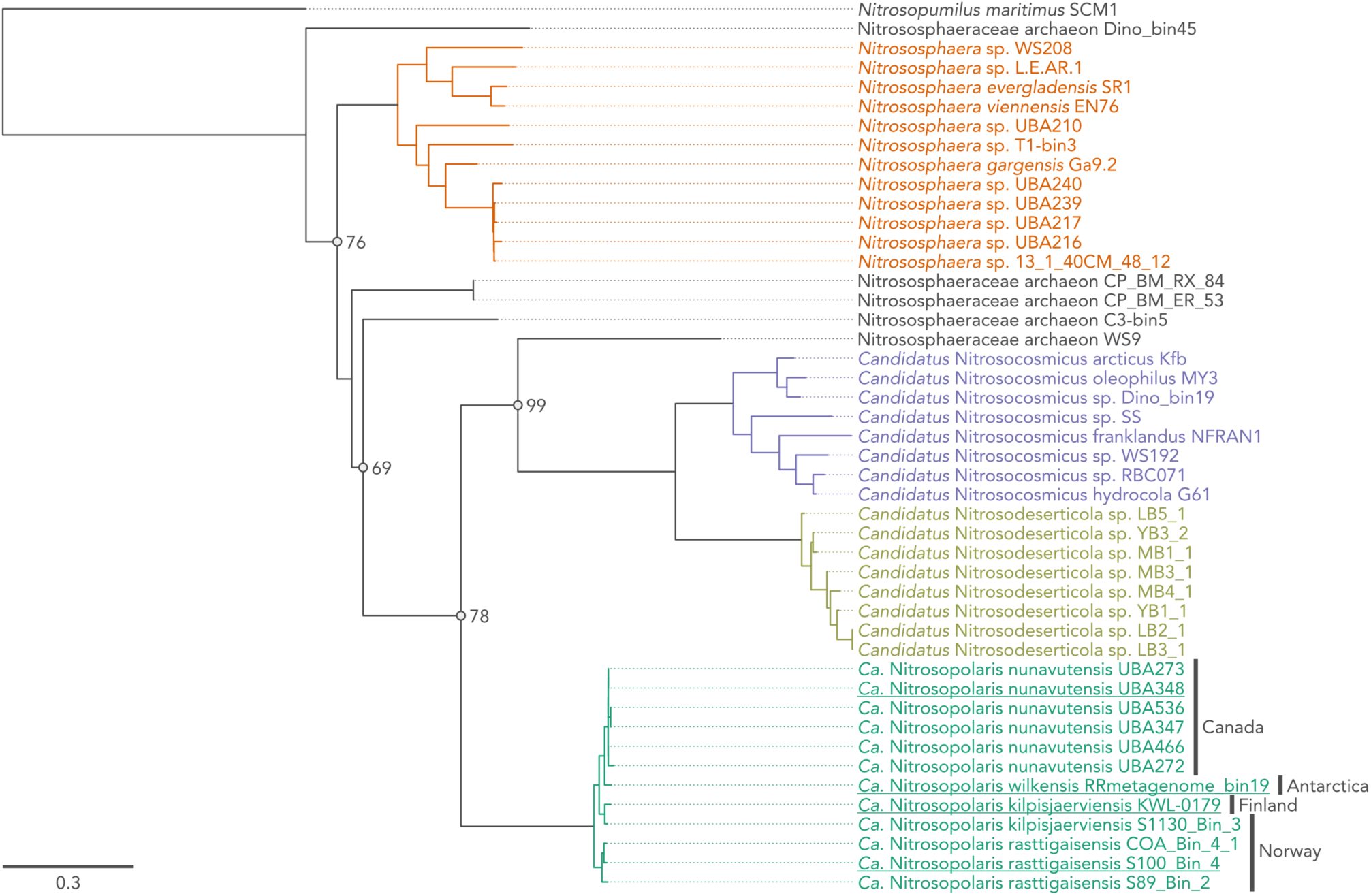
Phylogenomic analysis of the UBA10452 lineage (*Candidatus* Nitrosopolaris). Maximum likelihood tree based on 59 single-copy genes from 12 metagenome-assembled genomes (MAGs) assigned to the UBA10452 lineage and 33 other Nitrososphaeraceae genomes available on GenBank. *Nitrosopumilus maritimus* SCM1 was used for rooting the tree. Nodes are supported by bootstrap values of 100% unless shown otherwise. Representatives for the four proposed species are indicated in underscore. This is an uncollapsed and bootstrapped version of the tree found in **Fig. 2a**.

**Suppl. Figure S4.**
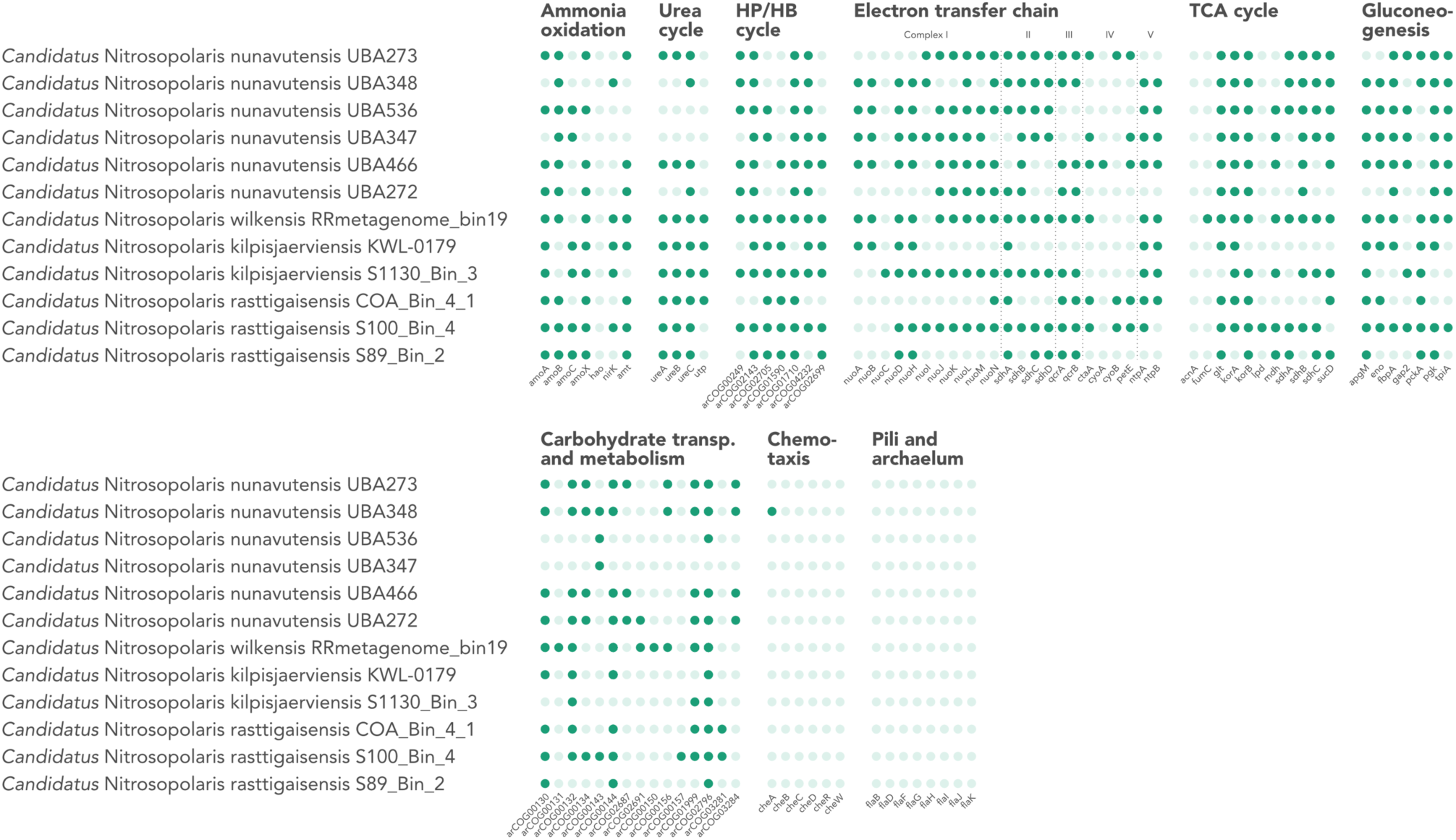
Metabolic potential of the UBA10452 lineage (*Candidatus* Nitrosopolaris). Metabolic potential was estimated based on the presence of key genes involved in selected pathways. Detailed information about the genes can be found in **Suppl. Table S2**.

**Suppl. Figure S5.**
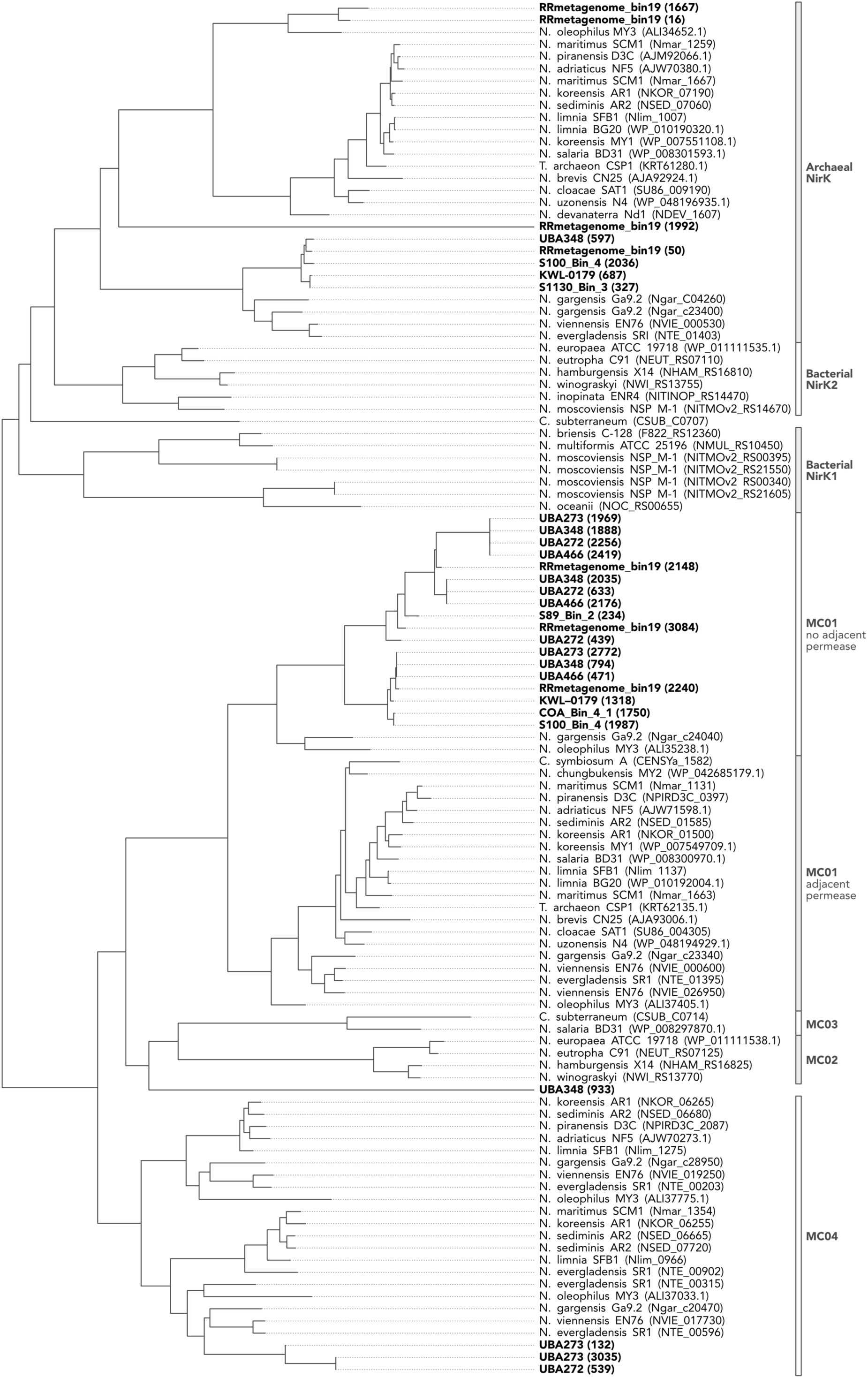
Phylogenetic analysis of putative NirK sequences from metagenome-assembled genomes (MAGs) belonging to the UBA10452 lineage (*Candidatus* Nitrosopolaris). Sequences from the UBA10452 are shown in bold and respective gene calls are given inside parenthesis. Other sequences were retrieved from Kerou *et al*. (2016).

